# Genomic LOCI Modulating Retinal Ganglion Cell Death Following Elevated IOP in the Mouse

**DOI:** 10.1101/212753

**Authors:** Felix L. Struebing, Rebecca King, Ying Li, Jessica N. Cooke Bailey, NEIGHBORHOOD consortium, Janey L. Wiggs, Eldon E. Geisert

## Abstract

The present study was designed to identify genomic loci modulating the susceptibility of retinal ganglion cells (RGC) to elevated intraocular pressure (IOP) in the BXD recombinant inbred mouse strain set. IOP was elevated by injecting magnetic microspheres into the anterior chamber and blocking the trabecular meshwork using a handheld magnet to impede drainage. The IOP was then measured over the next 21 days. Only animals with IOP greater than 25 mmHg for two consecutive days or an IOP above 30 mmHg on a single day after microsphere-injection were used in this study. On day 21, mice were sacrificed and the optic nerve was processed for histology. Axons were counted for both the injected and the control eye in 49 BXD strains, totaling 181 normal counts and 191 counts associated with elevated IOP. The axon loss for each strain was calculated and the data were entered into genenetwork.org. The average number of normal axons in the optic nerve across all strains was 54,788 ± 16% (SD), which dropped to 49,545 ± 20% in animals with artificially elevated IOP. Interval mapping demonstrated a relatively similar genome-wide map for both conditions with a suggestive Quantitative Trait Locus (QTL) on proximal Chromosome 3. When the relative axon loss was used to generate a genome-wide interval map, we identified one significant QTL (*p*<0.05) on Chromosome 18 between 53.6 and 57 Mb. Within this region, the best candidate gene for modulating axon loss was *Aldh7a1*. Immunohistochemistry demonstrated ALDH7A1 expression in mouse RGCs. *ALDH7A1* variants were not significantly associated with glaucoma in the NEIGHBORHOOD GWAS dataset, but this enzyme was identified as part of the butanoate pathway previously associated with glaucoma risk. Our results suggest that genomic background influences susceptibility to RGC degeneration and death in an inducible glaucoma model.

## Introduction

Glaucoma is a diverse set of diseases with heterogeneous phenotypic presentations. Untreated, glaucoma leads to permanent damage of axons in the optic nerve and visual field loss. Millions of people worldwide are affected (Quigley, 1996; Thylefors and Negrel, 1994), and in the United States glaucoma is the second leading cause of blindness (Leske, 1983). Adult-onset glaucoma is associated with multiple risk factors such as intraocular pressure (IOP), age, family history, ethnicity, central corneal thickness and axial length (Aboobakar et al., 2016; Liu and Allingham, 2011; Nickells, 2012; Springelkamp et al., 2017). Genetically, primary open angle glaucoma (POAG) and normal tension glaucoma (NTG), common forms of adult-onset glaucoma, are currently associated with 16 loci (Wiggs and Pasquale, 2017) and exhibit complex inheritance. The present set of POAG genetic risk factors explains only 3-6% of the genetic variance (Verma et al., 2016). Thus, fully defining the genetic basis of glaucoma will require advances in analytical approaches and experimental methods (Boyle et al., 2017).

Many investigators have turned to mouse models to identify causal events linking glaucomatous risk factors and the death of retinal ganglion cells (Struebing and Geisert, 2015). Compelled by the genetic simplicity that recombinant inbred (RI) mouse strains afford in mapping genotype to phenotype (Geisert et al., 2009), we designed a study investigating the variance of axon loss due to increased IOP in a large cohort of mice. For this purpose, we used the BXD RI strain set, which is particularly suited for the study of genetics and the effects on the severity of glaucoma. This genetic reference panel presently consists of over 200 strains. The original 36 strains were extensively studied by our group for more than 18 years. Over the last decade, the Williams group has greatly expanded the BXD family. Both parental strains (C57BL/6J and DBA/2J) are completely sequenced, and all BXD strains are fully mapped with more than 17,000 genetic markers. As a result, the current set of BXD strains is far larger than any other mature RI resource and allows for sub-megabase resolution QTL mapping. Furthermore, this strain set was previously used to develop several large data sets, including numerous ocular phenotypes (RGC number, IOP, eye size, retinal area, etc.), global transcriptome data for the eye (HEIMED (Geisert et al., 2009)) and retina (HEI Normal Retina Database (Freeman et al., 2011), DoD Normal Retina database (King et al., 2015)), and experimental consequences of optic nerve crush (ONC Retina Database (Templeton et al., 2013)) along with the effects of blast injury (DoD Blast Dataset (Struebing et al., 2017)). Each data set consists of measurements for as many as 80 strains, making the BXD set extremely revealing in the analysis of QTLs and ocular disease networks.

Because we needed a consistent and predictable model to increase IOP with little technical variation, we have adopted a method developed by Samsel et al. (Samsel et al., 2010) to induce acute IOP elevation through the injection of magnetic microspheres and the migration of these microspheres into the trabecular meshwork using magnetic fields. This method of blocking the trabecular meshwork yields a reliable IOP increase, and allows us to define BXD strains with differing susceptibility to axon loss. We then use these data to define a genomic locus modulating axon loss across the BXD strains.

## Methods

### Animals

All animals used in this study were either bought from the Jackson laboratory (Bar Harbor, ME) or bred in-house in a parasite-free facility. All mice were between 60 and 100 days of age and maintained on a 12h light – 12h dark cycle with food and water *ad libitum*. All procedures involving animals were approved by the Animal Care and Use Committee of Emory University and were in accordance with the ARVO Statement for the Use of Animals in Ophthalmic and Vision Research.

### Microsphere Injection and Elevation of IOP

To define the effect of elevated IOP on retinal ganglion cell loss, we examined axonal loss in 47 BXD RI strains. A total of 230 mice received an injection of magnetic microspheres into the right eye, following a method developed by Samsel et al. (Samsel et al., 2010). Briefly, magnetic microbeads (Spherotech PM-40-10) with a diameter of 4.14µm were washed in sterile HBSS 3 times and diluted 3-fold. The animals were deeply anesthetized with Ketamine (100mg/kg) and Xylazine (15mg/kg) and 10-15 µL of microbead solution were slowly injected into the anterior chamber with a custom-made 30G needle. The needle was held in place while microspheres were pulled into the trabecular meshwork with the help of a handheld 0.5T magnet. After having selectively fixed the magnetic microbeads in the iridocorneal angle, the needle was retracted and carrier HBSS was allowed to drain from the anterior chamber. After the end of the surgery, a drop of topical antibiotic (Certi-Sporyn Neomycin sulfate) was applied to the cornea and mice were allowed to recover on a heating pad until fully awake. Left eyes of many animals served as non-injected controls, but were excluded when any axon damage was present. A total of 181 control eyes and 191 bead injected eyes were included in the study.

### IOP Measurement

We measured IOP before the injection of magnetic microspheres to establish baseline values, and then at 2d, 4d, 7d, 9d, 14d, and 21d after the surgery. A rebound tonometer (Tonolab Colonial Medical Supply) was used to measure the IOP under anesthesia with 5% Isoflurane. Measured values are averaged values of 18 (3×6) repeated measurements per animal and time point. IOP readings obtained with the Tonolab instrument have been shown to be accurate and reproducible in various mouse strains, including DBA/2J (Nagaraju et al., 2007; Saleh et al., 2007). The visual axis of the injected eyes remained clear with no visible signs of inflammation. For a bead-injected eye to be included in the sample set, the IOP had to be above 25 mmHg in two consecutive measurements or above 30 mmHg in a single measurement. Bead injected eyes that did not meet this minimum criterion were excluded from the analysis.

### Analysis of Optic Nerve Axon Damage

After a survival period of 21 days, the mice were anesthetized with Ketamine (100mg/kg) and Xylazine (15mg/kg) and perfused through the heart with saline followed by 2% paraformaldehyde and 2% glutaraldehyde in phosphate buffer (pH 7.4). The optic nerves were carefully dissected and post-fixed for 24 hours at 4°C. Nerves were dehydrated and embedded in plastic (Libby et al., 2005). Thin sections (0.7µm) were cut from each nerve and mounted on glass slides. Sections were stained using a modified paraphenylenediamine (PPD) staining protocol. Slides were photographed using an Olympus BX51 microscope and a Microfire camera (Optronix) at a final magnification of 1,200X. Healthy axons were counted using ImagePad software developed in our laboratory for the iPad as described previously (Templeton et al., 2014). To assure that all sections were sampled systematically, a grid overlay was placed on the image of the optic nerve and fields were counted every 10 cells along appropriately spaced rows. This resulted in a minimum of 20 fields evenly spaced across the optic nerve, assuring that all sectors of the nerve were included in our sample. The sample was then converted to the average number of axons per mm^2^. The cross-sectional area of the section through the optic nerve was calculated from a low power photomicrograph using NIH ImageJ. These two measures were then used to provide an accurate estimate of the axons within each optic nerve.

### Analysis of Axon Loss

To map the genomic loci modulating axon loss, we calculated the average number of axons per strain in the normal nerves and the average number of axons in each strain following elevated IOP. We then subtracted the average number of axons in the bead injected nerves from that of the normal nerves to define the axon loss for each strain. Calculated values were used to map QTLs modulating the changes that occur following elevated IOP.

### Candidate Gene Selection Within the QTL Region

Our approach for identification of candidate genes involved the following steps: **1)** trait mapping and definition of the 95% confidence interval; **2)** haplotype analysis using our new sequence data to restrict analysis to those parts of the QTL that are not identical-by-descent between the parental strains; **3)** identification and analysis of all polymorphisms, including SNPs, InDels, inversions, and CNVs in the QTL interval using D2 genome sequence and the reference genome sequence of B6; **4)** identification of genes whose expression is high in the eye, retina, or other ocular tissue; **5)** correlation between our phenotype data and gene expression data from the eye and/or retina to extract highly correlated genes in the QTL; and **6)** gene analysis from the literature and other external data resources.

### Immunohistochemistry

For tissue preparation, mice were deeply anesthetized with Ketamine (100mg/kg) and Xylazine (15mg/kg) and perfused through the heart with saline followed by 4% paraformaldehyde in phosphate buffer (pH 7.3). The dissected eyeballs were post-fixed in 4% paraformaldehyde for an extra hour at room temperature. Eyeballs were incubated with 30% sucrose in phosphate buffered saline (PBS) at 4˚C overnight and embedded within Optimal Cutting Temperature compound (OCT). Cryosections, 12-15 µm thick, were cut on a Leica CM 1850 cryostat and kept in a −20°C freezer until further processing. For whole mounts, retinas were removed from the globe and cut into quarters. Sections or whole retinas were rinsed in PBS with 1% Triton-X100 and blocked with 5% normal donkey serum and 5% BSA in PBS with 1% Triton-X100, and placed in primary antibodies ALDH7A1 (Abcam, ab154264, Cambridge, MA) at 1:1000 and TUJ1 (gift from Anthony Frankfurter) at 1:1000 overnight at 4˚C. The sections or whole mount retinas were rinsed in PBS and then placed in secondary antibodies (Alexa Fluor 488 AffiniPure Donkey Anti-Rabbit, Jackson Immunoresearch, Cat. #715-545-152 and Alexa Fluor 594 AffiniPure Donkey Anti-Mouse, Jackson Immunoresearch, Cat. #715-585-150, West Grove, PA) at 1:1000 and To-PRO-3 Iodide (Molecular Probes, Cat. # T3605, Waltham, MA) at 1:1000 as nuclear counterstain for two hours at room temperature. After 3 washes in PBS, cover slips were placed over the sections or retina whole mounts using Fluoromount – G (Southern Biotech, Cat. # 0100-01, Birmingham, AL). High-resolution Z-stacks were captured on a Nikon confocal microscope with the Nikon C1 software. Z-stacks were collapsed using imageJ and the whole image was adjusted for contrast and brightness using Adobe Photoshop.

### QTL validation using qPCR

Retinas from 4 B6 and 4 D2 mice were dissected on ice into Hank’s Balanced Salt Solution (HBSS) with RiboLock Rnase inhibitor (Thermo Scientific). RNA was isolated using the RNeasy Mini Kit on a Qiacube (Qiagen) and including a DNase1 gDNA digestion step. First strand synthesis was carried out according to the manufacturer’s instructions (Takara PrimeScript Real Time). Primers were designed using NCBI Primer Blast and validated for specificity by melting curve analysis and Sanger sequencing of amplicons. The sequences were Aldh7a1 fwd (5’-GGAGCTGTATTTCCGGGGCT-3’) and Aldh7a1 rev (5’-CGCGTTTTGGGGCAGGAATA-3’), and the amplicon crossed an exon junction between exon 3 and 4 (NM_138600.4). Quantitative cycling was carried out on a realplex2 Mastercylcler (Eppendorf) for 40 cycles and 60° annealing temperature using SYBR Green qPCR Master Mix (Qiagen) following the manufacturer’s protocol. Reactions were run in quadruple and *Ppia* served as housekeeping gene (Qiagen QuantiTect Primer assay). Reported values are dCT values.

### Human glaucoma association analysis (NEIGHBORHOOD dataset)

The peak area of association on mouse chromosome 18 was examined for associations in human datasets, specifically looking for an association of ALDH7A1 with POAG. Liftover of mouse genomic coordinates resulted in one whole syntenic region located on human chromosome 5. This region was queried for associations with POAG in the NEIGHBORHOOD (Bailey et al., 2016) dataset.

## Results

The IOP was elevated in 49 strains of mice by injection of magnetic beads into the anterior chamber and fixation thereof in the trabecular meshwork with a hand-held magnet (Samsel et al., 2010). In all cases, the right eye was injected, while the left eye served as control except in a few cases where the left optic nerve contained damaged axons. Following the injection, IOP was measured in both eyes over the next 21 days. The IOP in the injected eye rose quickly to glaucomatous levels (typically defined by > 20mmHg in humans) and remained elevated for the duration of this experiment (Fig. 1). Thus, this method proved to be a reliable approach to acutely elevate IOP in the mouse eye, even though it was originally developed for the rat (Samsel et al., 2011).

**Figure 1.**
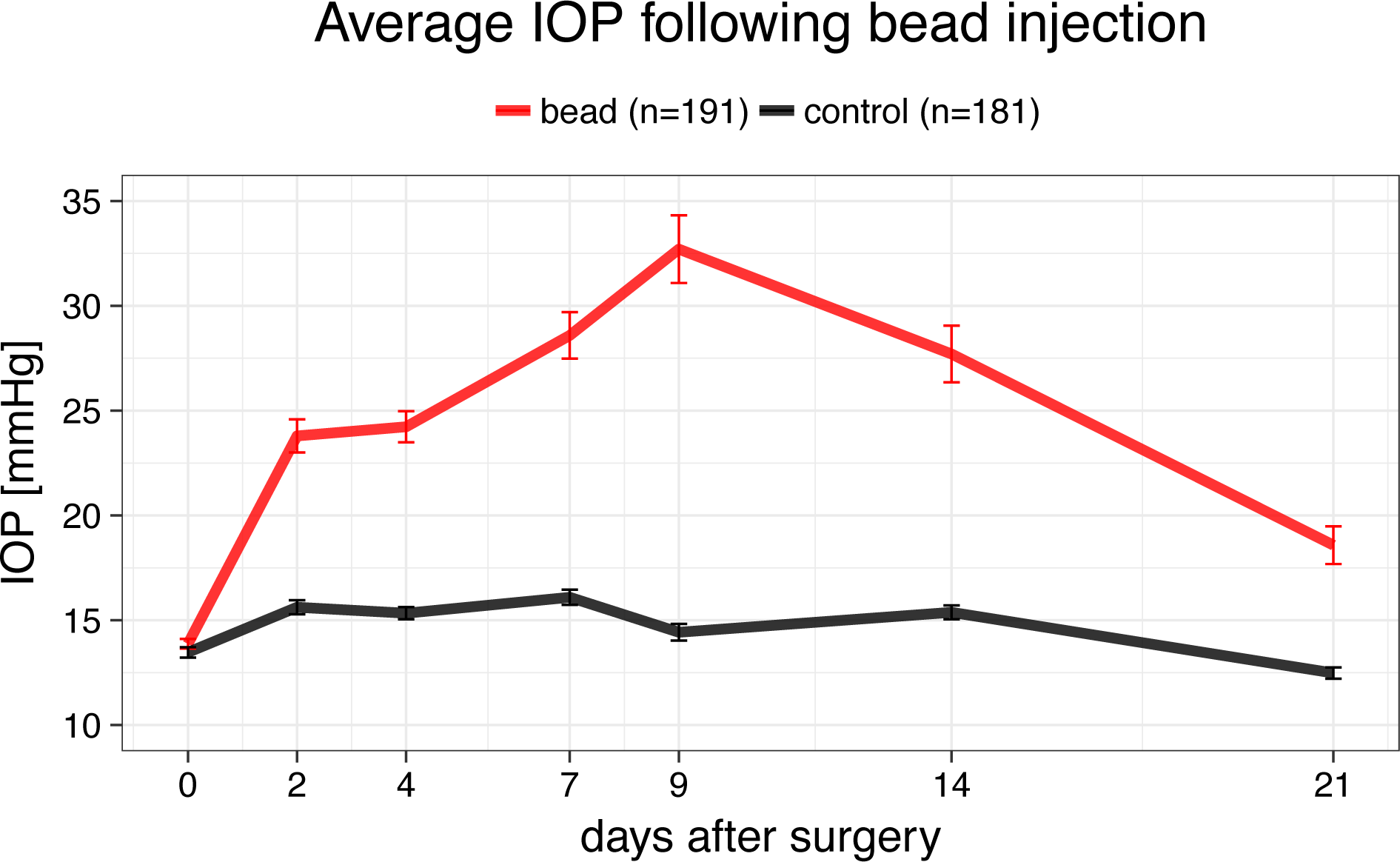
IOP development over 21 days after bead injection surgery. IOP was measured at the days given on the X-axis and averaged across all strains. IOP rises quickly on the second postoperative day, peaks at 9 days and stays elevated throughout the whole 3 weeks. Error bars show standard error.

The first step in the analysis was to generate a series of measures for the normal control eyes. After appropriate fixation and preparation of the optic nerve for microscopy, normal axons were quantified for 181 eyes in 49 strains. (Fig. 2, right panel). The number of axons ranged from a mean of 37,2277 axons in BXD36 to a mean of 78,159 in BXD97. The average number of axons per eye was 54,788 with a standard deviation of 8,622, demonstrating natural variation in the number of optic nerve axons across the BXD strain set. There was a significant difference between the strain with the highest and the lowest number of axons within the optic nerve (p = 0.002, Mann-Whitney U test).

**Figure 2.**
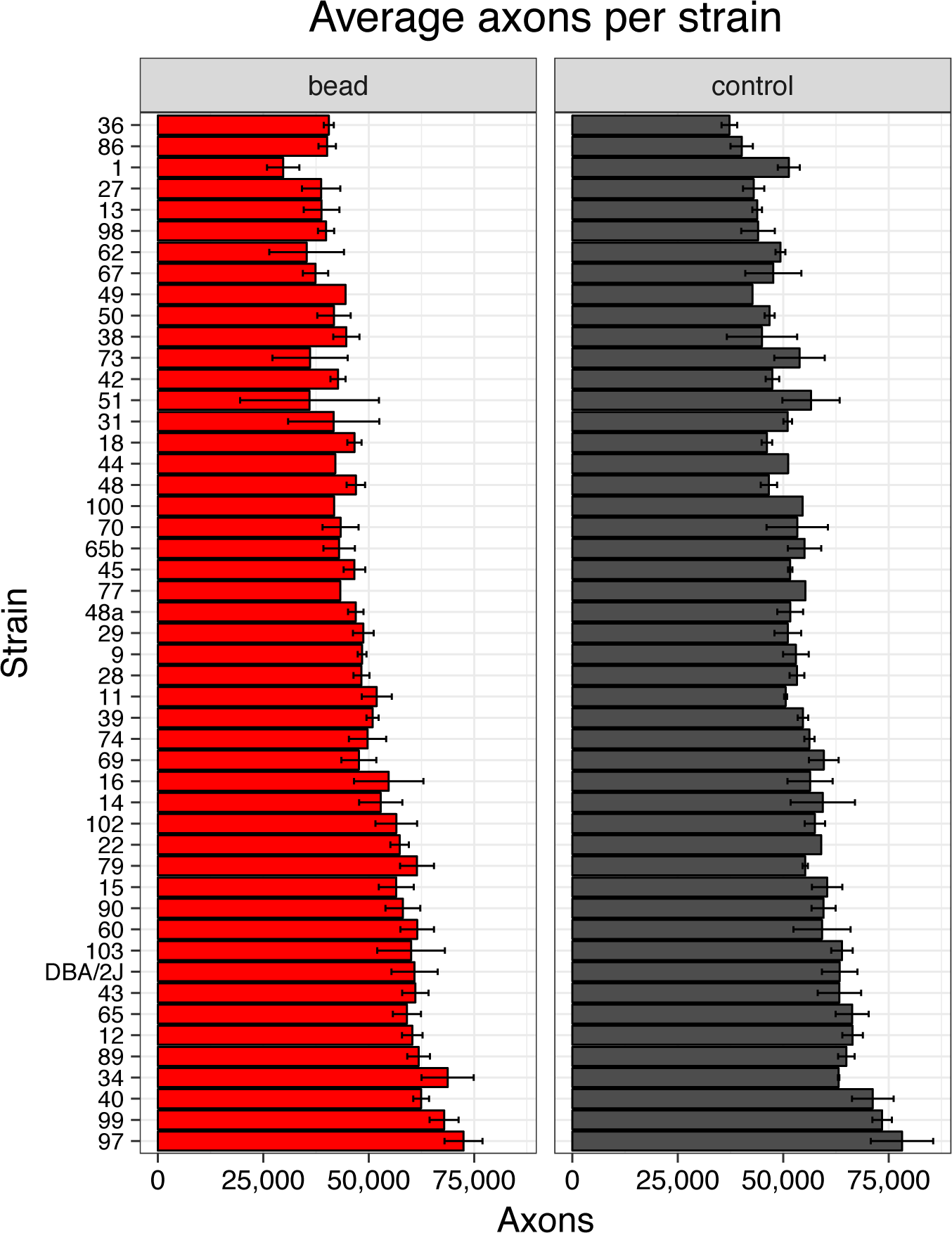
Axon counts for the normal eye (‘control’, grey) and for the eye 21 days after elevation of IOP (‘bead’, red). Error bars show standard error.

When the IOP was artificially elevated, the average axon number decreased across the 49 strains to 49,545 axons per optic nerve with a standard error of 9,982 axons (Fig. 2b). Thus, elevation of IOP decreased the average number of axons in the optic nerve, while it increased the standard deviation. Within the bead injected eyes, axon number ranged from 29,720 axons in the lowest strain (BXD1) to 72,473 axons in the highest strain (BXD97). The difference between the lowest and highest strain was significant (p=0.018, Mann-Whitney U test). When the number of axons in the normal retina and the number of axons following elevated IOP were used to generate genome-wide interval maps, they both revealed the same suggestive quantitative trait loci (QTLs) on proximal chromosome 3 (Fig. 3). Neither dataset had QTLs that reached the significance level (p > 0.05).

**Figure 3.**
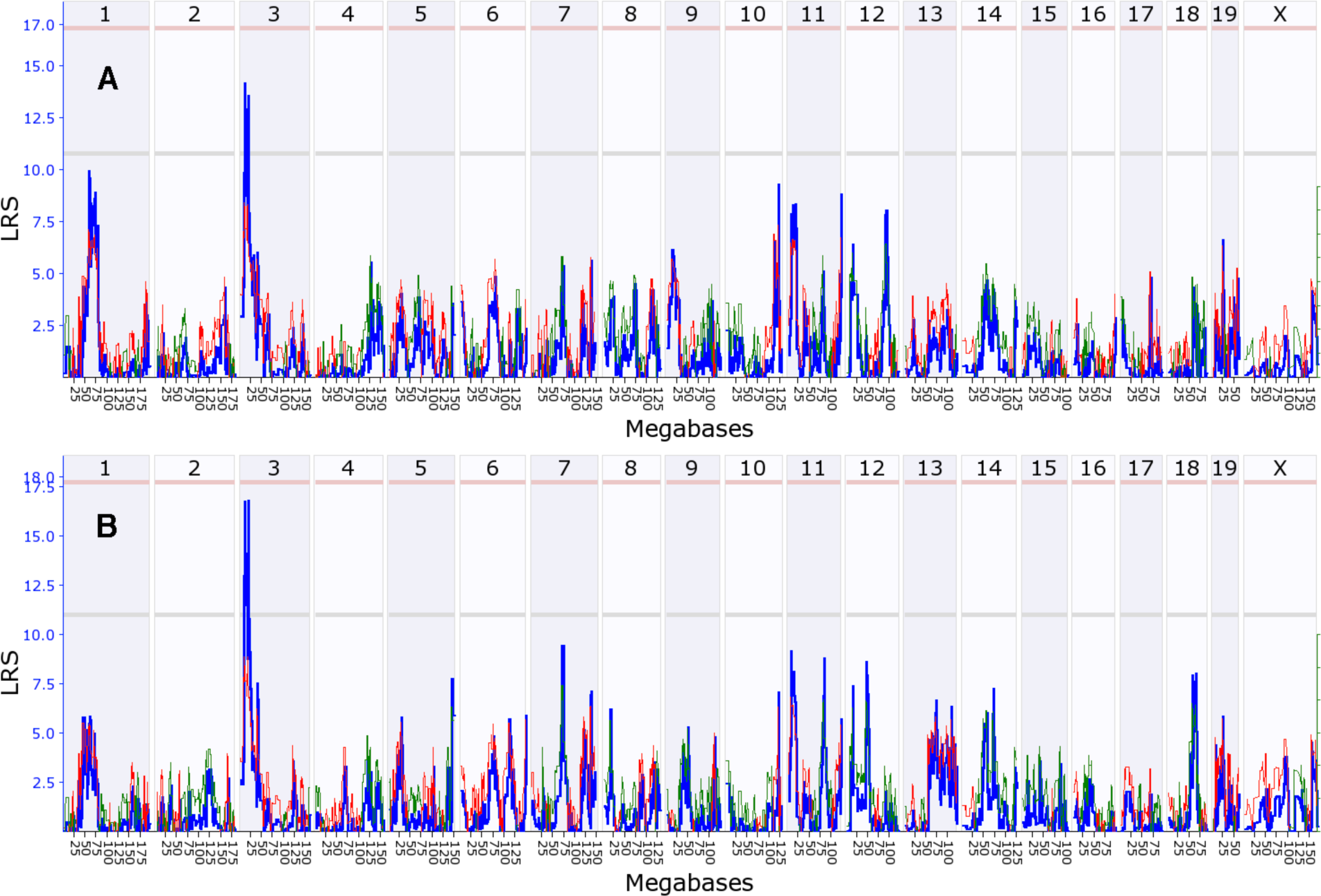
Interval map of axon number across the mouse genome is illustrated for the normal eye (A) and the eye 21 days following elevation of IOP (B). The blue line indicates the total likelihood ratio statistic (LRS) score as specified on the y-axis. The red line illustrates the contribution from the B6 allele and the green line the contribution from the D2 allele. The location across the genome is indicated on the top from chromosome 1 to chromosome X. Notice one suggestive QTL peak on Chr3 (exceeding the gray line, p<0.63).

To define genomic loci that could modulate the susceptibility of RGCs to death, axon loss per strain was calculated by subtracting the mean number of axons in the bead injected eyes from the mean number of axons in the normal eyes for each strain (Fig. 4). The eye with elevated IOP had an average axon loss of 9.6% relative to the non-injected control eye. There was considerable variability in axon loss across the 49 strains ranging from basically no axon loss in several strains to an average loss of 20,000 axons in BXD1 and BXD51. These data were used to generate a genome-wide interval map (Fig. 5), which revealed a suggestive QTL on chromosome 9 and a significant QTL on chromosome 18. The significant peak on chromosome 18 (Fig. 6) was found between two genomic markers, rs13483369 (54.614840 Mb, mm10) and rs215803889 (56.037647 Mb). Because the high BXD genotyping density with >17,000 segregating markers allows us to map with a precision of roughly ±1 Mb, we examined the region on chromosome 18 from 53.6 to 57 Mb to define potential candidate genes modulating axonal loss. Good candidates should either contain nonsynonymous SNPs altering their amino acid sequence, or they should be cis-eQTLs, meaning that their expression level segregates into two groups defined by the distribution of parental alleles (Struebing et al., 2016). Examining the sequences between B6 and D2 using the SNP browser on GeneNetwork did not reveal genes with nonsynonymous SNPs within this 2 Mb large region. We then queried our recently published BXD Normal Retina databases on GeneNetwork for cis-eQTLs (DoD Normal Retina Affy MoGene 2.0 ST (May 15) RMA Gene Level, (King et al., 2015)). Only one known gene, *Aldh7a1*, showed a significant LRS score (p < 0.01, Probe 17354434), suggesting that this gene is a single cis-eQTL within this interval. The differential expression between C57BL/6J and DBA/2J parents was then confirmed by quantitative PCR (p = 0.028, Wilcoxon rank-sum test, Supplemental Fig. 1).

**Figure 4:**
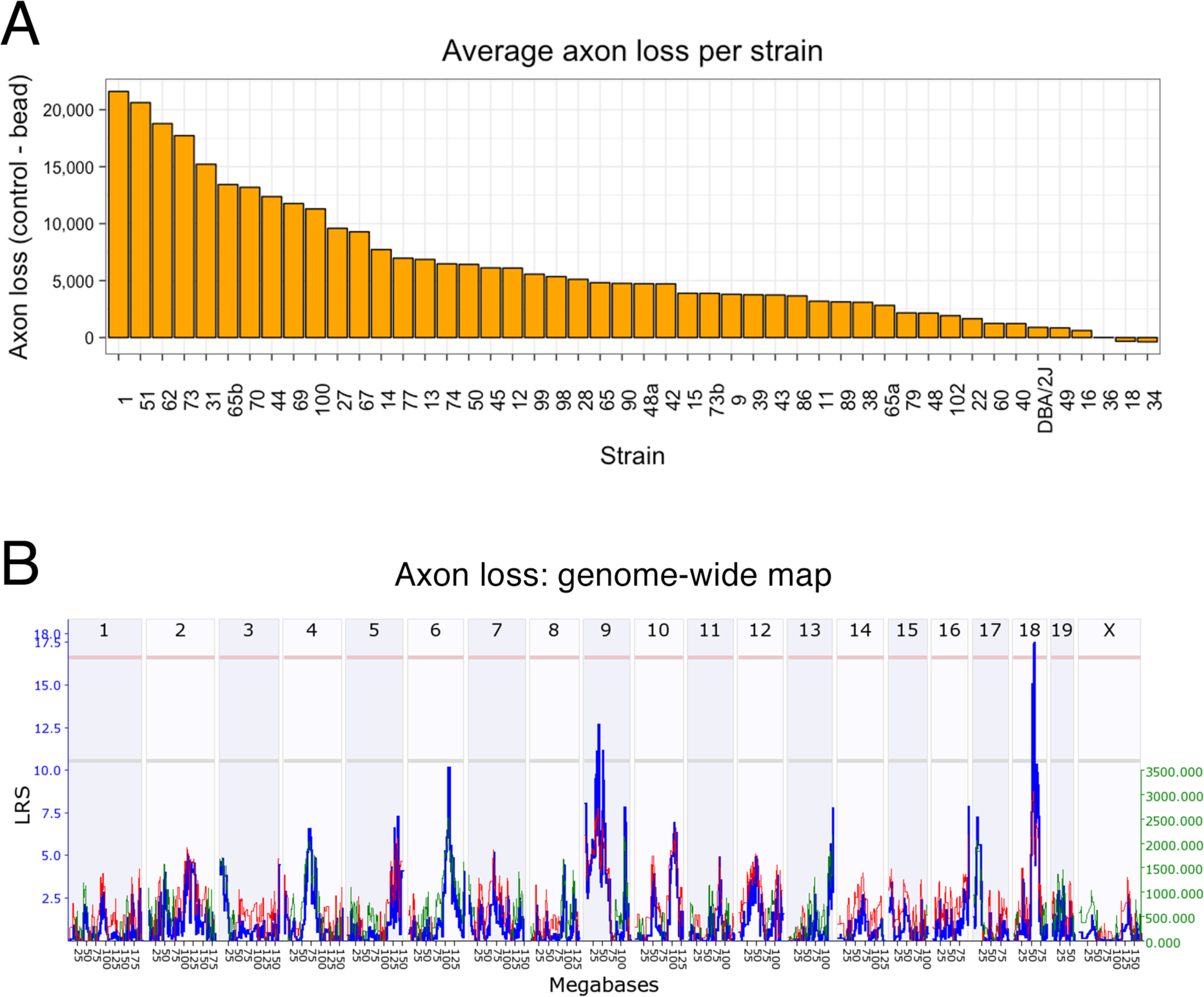
The average axon loss per strain is given in (A) in decreasing order. The genome-wide interval map is shown in (B). Here, chromosomes are displayed on the X axis, while the LRS score is plotted in blue along the Y axis. The light red line defines the genome-wide QTL significance interval of p < 0.05, while the light grey line identifies a suggestive QTL. Notice one significant LRS peak on chromosome 18 and one suggestive QTL on chromosome 9.

**Figure 5.**
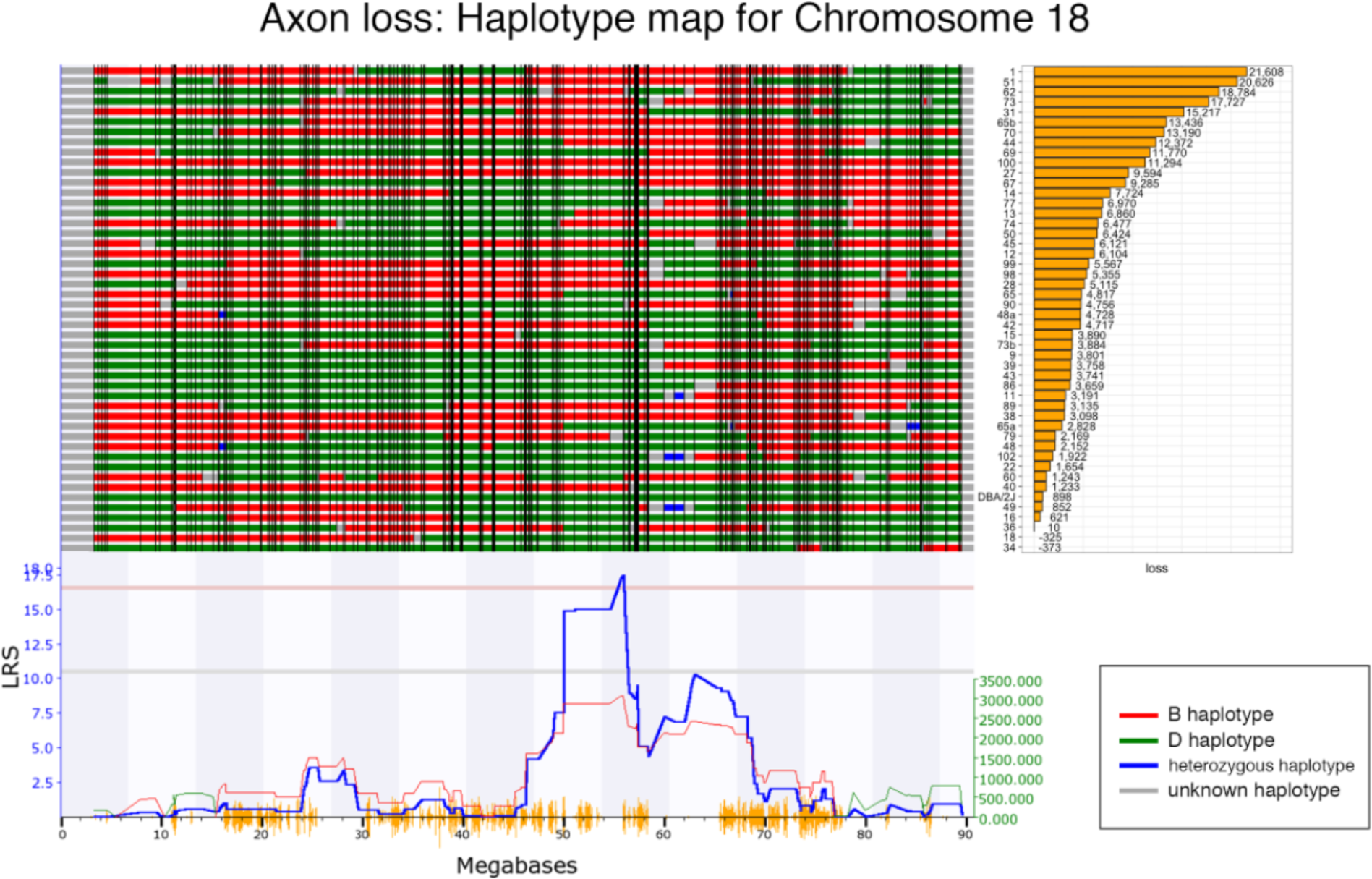
Map of gene locations across Chr. 18. The haplotype map for the 47 strains in the axon loss dataset is shown in the top panel. The haplotype map displays the contribution of each of the two parental strains. The red represents the B6 alleles, green defines the D2 alleles, blue represents regions of the DNA that are heterozygotic and gray is unmapped. At the far right is a list of the specific BXD RI strains with the associated axon loss in decreasing order from top to bottom. The QTL significance is indicated by the blueline below the haplotype map. A positive additive coefficient (red line) indicates that B6 alleles are associated with higher trait values. The QTL reaches significance from ~56 to ~58 Mb (p<0.05, light red line).

**Figure 6.**
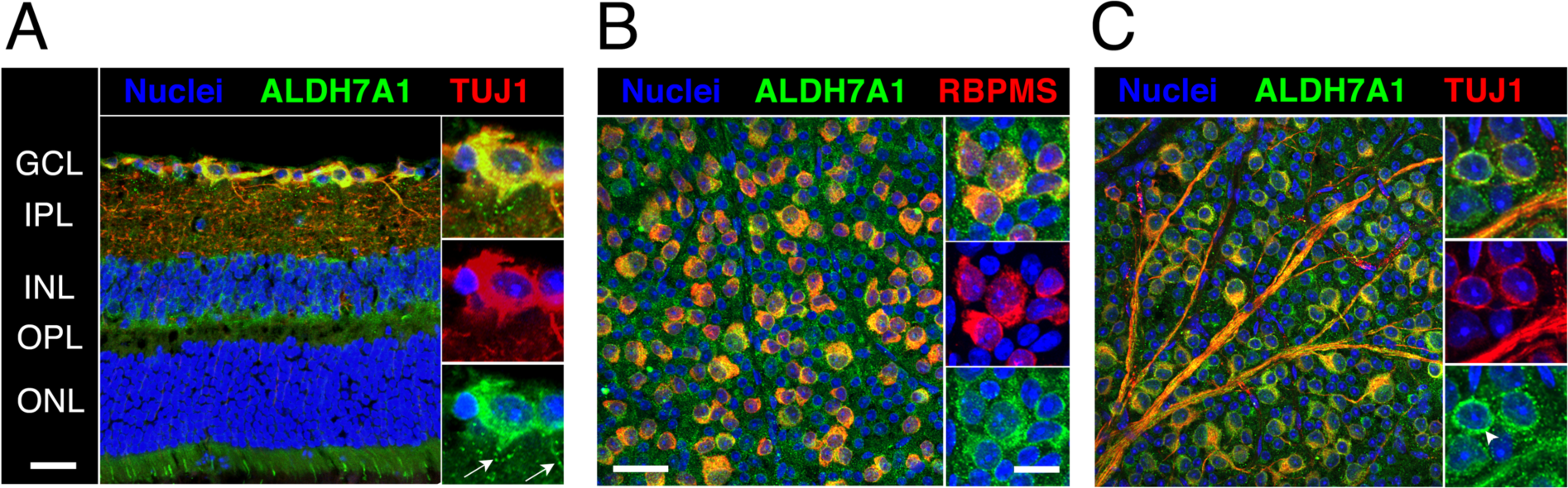
Cross section of a C57BL/6J retina stained for nuclei (blue), ALDH7A1 (green), and the neuronal marker TUJ1 (Class III beta-tubulin, red) is shown in (A). Notice the colocalization of ALDH7A1 with RGCs and its stippled distribution in RGC dendrites (white arrows, magnified section). In (B), a flat-mounted retina was stained with ALDH7A1 and RGC marker RBPMS and the ganglion cell layer (GCL) was imaged *en face*. While all RGCs are double labeled, some cells in the GCL stained exclusively for ALDH7A1 albeit in a less intense manner. (C) is analog to (B) except that it was stained for TUJ1, which also stains RGC axons. ALDH7A1 is most prominently distributed in a perinuclear fashion (arrowhead, magnified section). ALDH7A1 staining is absent in the nucleus, but slightly present in RGC axons. GCL = ganglion cell layer, IPL = inner plexiform layer, INL = inner nuclear layer, OPL = outer plexiform layer, ONL = outer nuclear layer. Low magnification scale bars = 50µm. High magnification scale bars = 20µm.

We then examined the distribution of ALDH7A1 in the retina using immunohistochemical methods. We stained retinal sections or flat-mounts with antibodies against ALDH7A1 and the RGC markers TUJ1 or RBPMS (Figure 6). While TUJ1 is known to stain RGCs and a few other retinal neurons as well as axons, RBPMS expression was shown to be restricted to the RGC population (Rodriguez et al., 2014; Struebing et al., 2016). ALDH7A1 co-localized with all TUJ1- and RBPMS-positive cells, while ALDH7A1 staining was also present throughout other retinal layers, including cone photoreceptors. Higher magnifications revealed that ALDH7A1 staining was relatively ubiquitous in cell body and axons but absent from the nucleus. These results are in line with other studies that have demonstrated mitochondrial and cytosolic localization of ALDH7A1 in humans and rodents (Brocker et al., 2011; Brocker et al., 2010; Wong et al., 2010).

In order to test whether variants in *ALDH7A1* influence primary open-angle glaucoma (POAG) susceptibility in humans, we interrogated results from the NEIGHBORHOOD GWAS glaucoma dataset from 3,853 cases and 33,480 controls in the human genomic region corresponding to the mouse *ALDH7A1* locus. The intronic variant rs62391530 showed the strongest relation to POAG with a nominal p-value of 0.02 and an effect size of −0.16. Even though ALDH7A1 variants did not reach the genome-wide significance threshold of 5×10^−8^, this gene is part of the KEGG “butanoate metabolic pathway”, which was previously identified to be associated with both POAG and NTG (Bailey et al., 2014).

## Discussion

Earlier work in glaucoma suggested a role for genetic background in the degeneration of RGCs. The best evidence for this comes from studies in inbred mice (Li et al., 2007). For example, Li et al. looked at the loss of retinal ganglion cells after optic nerve crush in 15 different inbred mouse strains. There was a considerable variability in ganglion cell loss among different strains, and the authors showed that this difference was heritable. In another study, Anderson et al. found that transferring the two known glaucoma-causing DBA/2J mutations onto a C57BL/6J wild-type background recapitulated the iris pigment disease without causing IOP elevation or a neurodegenerative optic nerve phenotype (Anderson et al., 2006). These studies clearly demonstrate a heritable component in the susceptibility of RGCs to cell death.

Here, we have shown that susceptibility to IOP-elevation induced RGC death is influenced by genetic background in the BXD RI mouse strain set. When we performed quantitative trait mapping for this trait, we could identify a significant genomic interval on Chromosome 18, which likely modified axon loss. We only found one candidate gene within this locus, and we were able to confirm differential expression of this gene, *Aldh7a1*, between the B6 and D2 parents. Furthermore, immunohistochemistry demonstrated expression of ALDH7A1 protein in RGC somata and axons, consistent with its hypothesized role of influencing RGC death susceptibility.

Aldehyde dehydrogenase 7 family member 1 (ALDH7A1), also known as antiquitin, is an enzyme involved in lysine metabolism and protection of cells from hyperosmotic stress (Brocker et al., 2011; Brocker et al., 2010). Mutations in *Aldh7a1* are most prominently known to cause pyridoxine-dependent seizures, a very rare neurological disease characterized by intractable pre- and neonatal seizures that can be treated with large daily doses of Vitamin B6 (Mills et al., 2006). A lower concentration of ALDH7A1 mRNA and protein was also previously found in the brain of DBA/2J mice compared to C57Bl/6J mice, which is coherent with our results and suggests that this QTL is not only retina-specific (Bhave et al., 2006).

While GWAS have identified many genes associated with glaucoma risk, a recent analysis (Bailey et al., 2014) used a randomized analysis to take the single gene data and assign associated molecular pathways. One pathway involved butanoate metabolism (KEGG pathway hsa00650; see Figure 3 in (Bailey et al., 2014)). This pathway is associated with both POAG (8 genes with nominal association) and NTG (8 genes with nominal associations) Of the 5 enzymes (ALDH1B1, ALDH2, ALDH3A2 ALDH7A1 and ALDH9A1) associated with the conversion of Acetaldehyde to Acetyl-CoA, three (ALDH1B1, ALDH2 and ALDH3A2) have a nominal association with either POAG or NTG. Even though ALDH7A1 does not have a significant association with human glaucoma, it is a participant in a molecular pathway that is directly involved in POAG and NTG risk.

Since we used an inducible glaucoma model in this study, additional factors may have influenced our mapping results. For example, the scleral stiffness (compliance) may be different between BXD strains, which could lead to differential susceptibility to artificial IOP elevation (Sigal and Ethier, 2009). Likewise, differences may exist in the clearance of magnetic beads from the iridocorneal angle. An interesting possibility for the difference in axon loss we have demonstrated in the BXD strain set is differential susceptibility of RGC subtypes to IOP elevation. It was previously shown that alpha-RGCs and melanopsin-containing RGCs were more resistant to axotomy-induced cell death (Duan et al., 2015; Struebing et al., 2016), and similar observations were made in a mouse model of chronically elevated IOP (Feng et al., 2013). These findings argue for the existence of distinct transcriptional programs within each RGC subtype (Struebing et al., 2016). Further developments in this area of research will certainly shed more light onto the molecular mechanisms leading to differential susceptibility of RGCs to death.

In summary, the present study examined the loss of RGCs following an elevation of IOP in the BXD mouse strain set. We identified a single locus on Chr. 18 that modulates the loss of RGCs. A good candidate gene within this locus is *Aldh7a1*. This gene may be important for RGC survival in the BXD mouse strains, and the ALDH7A1 enzymatic pathway is associated with both NTG and POAG (Bailey et al., 2014). These results underline how mouse models and systems genetics could be used to enhance the interpretability of human GWA studies.

## Acknowledgements

We would like to thank XiangDi Wang (Department of Ophthalmology, Hamilton Eye Institute) for her assistance in collecting data.

## Disclosures

This study was supported by an Unrestricted Grand from Research to Prevent Blindness, NEI grant R01EY178841 (E.E.G.), Owens Family Glaucoma Research Fund, P30EY06360 (Emory Vision Core, to PM Iuvone) R01 EY022305 (JLW), PO30EY014104 (JLW),

---

**Supplemental Figure 1.**
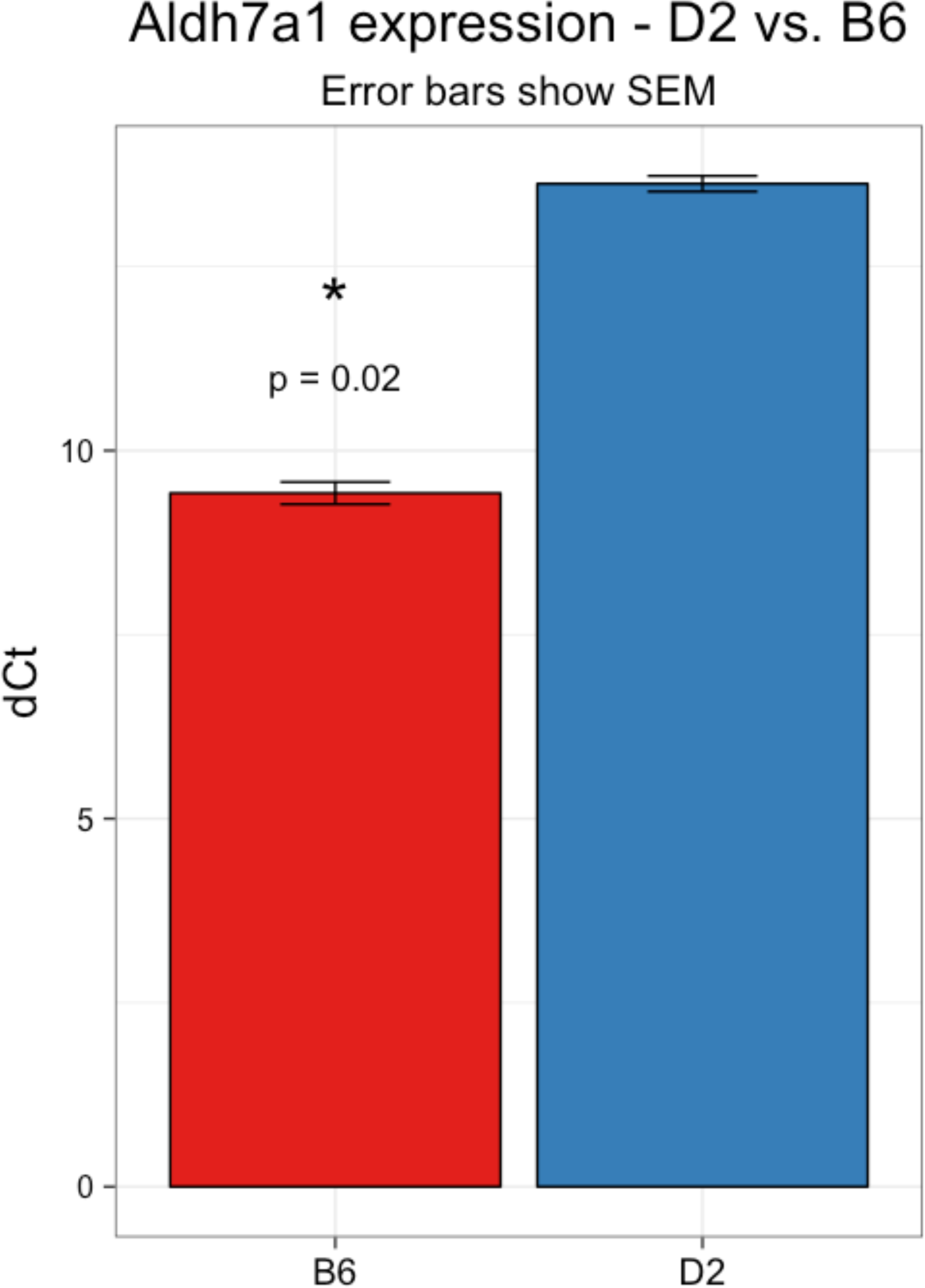
Aldh7a1 expression measured by qPCR in D2 vs. B6 mice. The Y-scale shows dCt values on a log_2_ scale after normalization with the housekeeping gene Ppia. Expression is significantly higher in retinas of B6 mice (p = 0.028, Wilcoxon rank-sum test).

